# Ligify 2.0: A web server for predicted small molecule biosensors

**DOI:** 10.1101/2025.10.20.683484

**Authors:** Simon d’Oelsnitz, Nicole Zhao, Pranay Talla, Jio Jeong, Joshua D. Love, Michael Springer, Pamela A. Silver

## Abstract

Prokaryotic transcription factors (TFs) are used as small molecule biosensors with broad applications in biotechnology, yet only a small fraction from microbial genomes have been characterized. To address this gap, we recently described the bioinformatic method Ligify, which leverages information from genome context and enzyme reaction databases to predict a TF’s cognate effector molecule. Here we report Ligify 2.0, a modern web server for Ligify predictions. We systematically evaluate 10,965 small molecules within the Rhea enzyme reaction database for associations to TFs, ultimately generating 13,435 hypothetical interactions between 1,362 small molecules and 3,164 TFs. We then develop an interactive web server (https://ligify.groov.bio) to search and visualize prediction data. Each TF sensor page includes visualizations for chemical ligand structures, interactive TF protein structures, and genome context. Pages also include metadata links, predicted promoter sequences, prediction confidence metrics, and references to relevant literature. A plasmid builder tool enables users to generate custom biosensor circuit designs. Finally, we provide case studies using Ligify 2.0 to identify two TFs from the pathogens *Escherichia coli* O157:H7 and *Mycobacterium abscessus* responsive to 4-hydroxybenzoate and Pseudomonas Quinolone Signal, respectively. The Ligify web server aims to facilitate the systematic characterization of biosensors for chemical-control of biological systems.

**Key points:**

- Ligify 2.0 contains >13,000 predicted transcription factor-small molecule interactions
- A rich web interface provides interactive visualizations and a plasmid design tool
- Predicted ligands for regulators from pathogenic bacteria are experimentally validated

**Graphic abstract:** 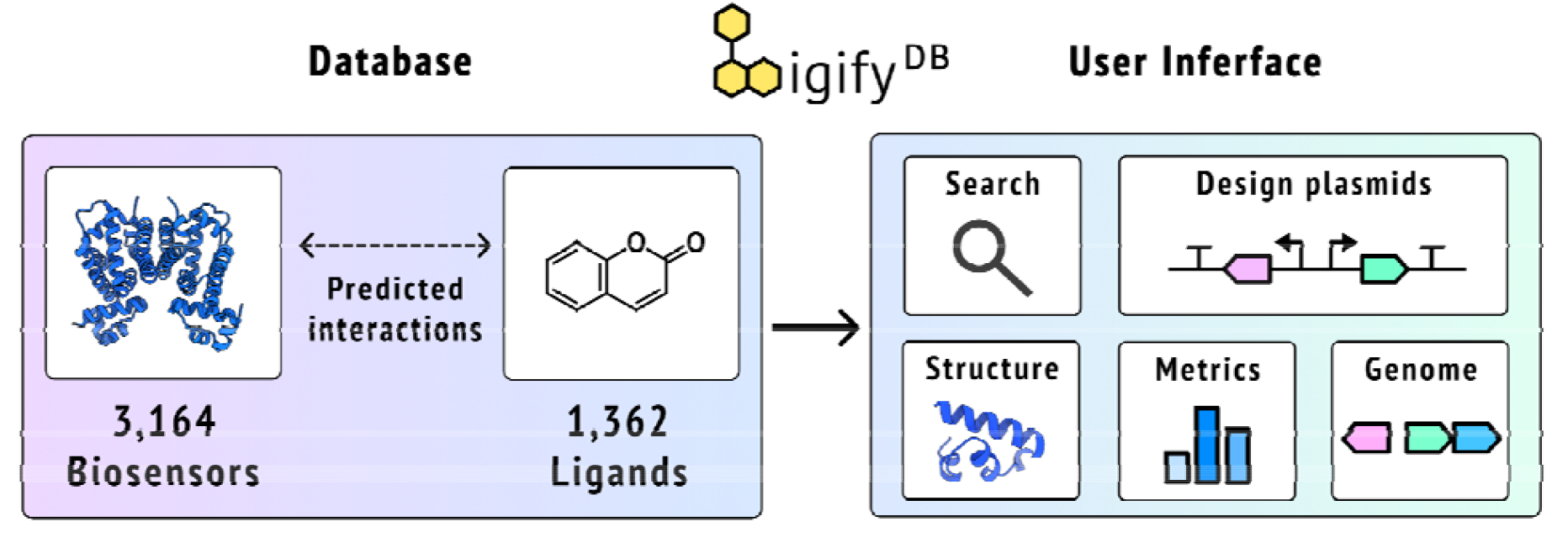

## Introduction

Genetically-encoded small molecule biosensors are core to many applications in biotechnology, ranging from high-throughput screening, diagnostics, responsive materials, and live-cell metabolite tracking^1–4^. Among biosensor modalities, prokaryotic ligand-inducible transcription factors (TFs) are one of the most widely used due to their broad host compatibility, compact size, high sensitivity, and wide dynamic range^5^. TFs regulate gene expression by undergoing ligand-induced conformational changes that modulate their DNA-binding affinity, thereby either promoting RNA polymerase recruitment or blocking its access to the promoter.

Despite their acknowledged utility, identifying a TF with desired ligand specificity remains challenging. Large genome modeling efforts have driven the characterization of TFs in model organisms, producing databases such as RegulonDB, BioCyc, and CoryneRegNet^6–8^. However, ligands for the vast majority of TFs outside these few organisms are still largely unknown. To help close this gap, genome context has proven to be effective at guiding the prediction of effector ligands for a given TF^9^. For example, a TetR-family regulator co-transcribed with an enzyme that glycosylates the antibiotic kijanimicin was found to bind specifically to kijanimicin as a ligand^10^.

Using genome context alone, recent software tools have automated the prediction of TF-ligand interactions. The Ligify and TFBMiner tools perform this prediction task by leveraging data from the enzyme reaction databases Rhea and KEGG, respectively, leveraging the phenomena that TFs sometimes recognize the substrates and/or products of adjacently encoded enzymes^11–14^. These tools aim to expand the knowledgebase of TF-ligand interactions contained within databases, such as groov^DB^, SensBio, RegPrecise, and the detectable molecules collection^15–18^. While useful, the tools suffer from several issues limiting their potential. All predictions must be computed on demand, which is slow, computationally expensive, limits use for bioinformaticians, and complicates a comprehensive analysis of prediction performance. Additionally, user interfaces are limited to the command line or simple text-based web applications, which delivers a poor user interface and hinders data interpretation.

Here, we report Ligify 2.0 (https://ligify.groov.bio), an interactive and extensible platform for predicted TF biosensors that overcomes accessibility, performance, and usability issues with existing TF prediction tools. Generated by running the Ligify workflow on over 10,000 chemicals in the Rhea database, Ligify 2.0 contains 3,164 unique TFs predicted to respond to 1,362 unique small molecules. Analyses indicate Ligify predictions cover a wide distribution of TFs from phylogenetically diverse hosts, and include a range of metabolites such as sugars, metals, amino acids, and aromatics. We then create an interactive interface to easily query predictions via text search, chemical similarity search, and attribute filtering, and visualize TF entry properties, such as their 3D protein structure, 2D chemical ligand structures, operons, predicted promoters, and prediction quality metrics. A plasmid designer tool is described, which facilitates the creation of reporter plasmids for experimental validation of predictions. Benchmarking results suggest Ligify can identify 32% of the validated sensor interactions in the groov^DB^ database. Finally, we experimentally validate predicted ligands for regulators from the bacterial pathogens *Escherichia coli* O157:H7 and *Mycobacterium abscessus*.

## Results

### Creating the database

We sought to improve the accessibility and analysis of TF biosensor predictions by pre-computing all possible results using Ligify. As described in the original publication, the Ligify workflow starts by using a small molecule input to fetch associated enzyme reactions from the Rhea database^14^ (**Figure 1)**.

**Figure 1:**
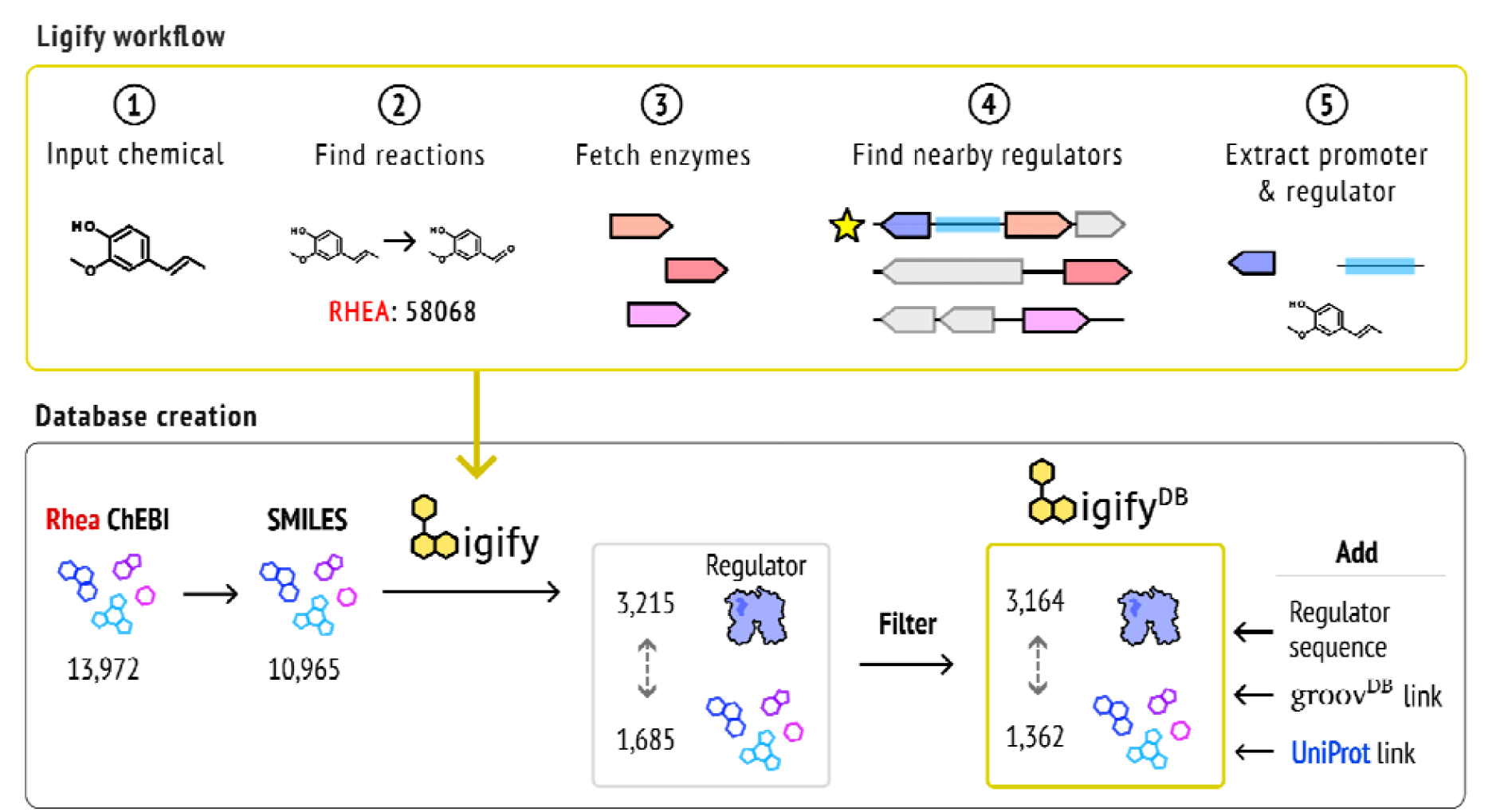
Workflow for creating the Ligify database. (top) The Ligify workflow, which takes a small molecule input and returns transcription factors predicted to interact with the molecule. (bottom) Workflow for creating Ligify^DB^. Ligify was run on all small molecule SMILES identifiers from Rhea, generating predictions for 1,685 molecules to 3,215 transcription factors. After filtering and data supplementation, the final version of Ligify^DB^ contains 13,435 predicted interactions.

Bacterial enzymes annotated to perform these reactions are fetched from Uniprot, and highly similar enzymes are filtered to reduce redundancy. Genes encoded within each enzyme’s operon are retrieved from the NCBI Gene Database, and their annotations are scanned for the terms “regulator”, “repressor”, and “activator” to identify TFs. If a TF is found, the corresponding promoter element is extracted, under the general assumption that TFs tend to regulate their own expression. If TFs are identified, the output of the Ligify workflow includes the TF gene and promoter, genes within the operon, associated literature, a rank calculated based on a set of heuristics, metadata links, and all possible candidate ligands.

To create a database containing all possible predictions from Ligify, we started by fetching all unique small molecules from the Rhea database in ChEBI format, which was 13,972 at the time of database generation. Each molecule was then converted into a SMILES format compatible with Ligify, which returned 10,965 molecules since some ChEBI entries did not have an associated SMILES identifier. After filtering out generic molecules, such as “water” or “hydron”, the Ligify workflow was performed on each molecule using default parameters (**see Methods**). Ultimately, 1,685 unique molecules were predicted to be associated with a total of 3,215 unique TFs. To reduce false positives we performed two filtering steps. First, we removed TFs smaller than 80 amino acids, since some of the shortest TFs known to bind DNA and a small molecule are around 90 amino acids long^19,20^. Next, we removed common co-factors, such as ATP and Coenzyme A, since these are overrepresented and unlikely to be the true ligands of any given TF (**see Methods**). Ultimately, this yielded 3,164 unique TFs associated with 1,362 unique molecules. To finalize the database, we (1) removed unnecessary data fields, such as operon sequences, to reduce size, (2) added the protein regulator sequence, (3) added Uniprot links to each regulator when possible, and (4) added links to homologous characterized TFs in the groov^DB^ database^18^. Ultimately, the final database is 12 megabytes and contains 13,435 hypothetical interactions.

### Analysis of Ligify content

To better understand the content of the resulting Ligify database, we performed a series of analyses. First we compared the diversity of organisms represented in Ligify^DB^. Among the 3,164 TF regulators, just over half belong to the Pseudomonadota phyla, with the remaining being represented by 29 other phyla dominated by Actinomycetota and Bacillota (**Figure 2a**). A closer look into the most common genuses reveals a prevalence of model organisms, such as Escherichia and Pseudomonas, as expected (**Figure 2b**). However, over 70% of regulators are not found within the top 10 organisms. Collectively, these results highlight the phylogenetic diversity of regulators within Ligify^DB^. We next analyzed the composition and distribution of regulator sequences, starting with structural family annotations. The most common structural families include LysR, TetR, GntR, and MarR families, which is expected given the wide representation of these families in nature^21^ (**Figure 2c**). Many less common families are also found in the database, such as the RpiR, PyrR, and PmpR structural families. Regulators tend to be between 150 and 350 amino acids long, with peaks around lengths of 180, 230, and 320, which typically correspond to the (MarR/TetR), LysR, and LacI families, respectively (**Figure 2d**). Finally, we compared the diversity of all 1,362 ligands within Ligify^DB^. Plotting the distribution of chemical structures using molecular fingerprints revealed a wide spread in chemical diversity, with larger clusters around sugars, aromatics, and amino acids, and smaller clusters around Coenzyme-A ligated molecules and cofactor-like molecules (**Figure 2e**). Among these, the most common ligands consisted largely of amino acids and central metabolites like pyruvate, likely because they can serve roles as general cofactors in enzymatic reactions^22^ (**Figure 2f**).

**Figure 2:**
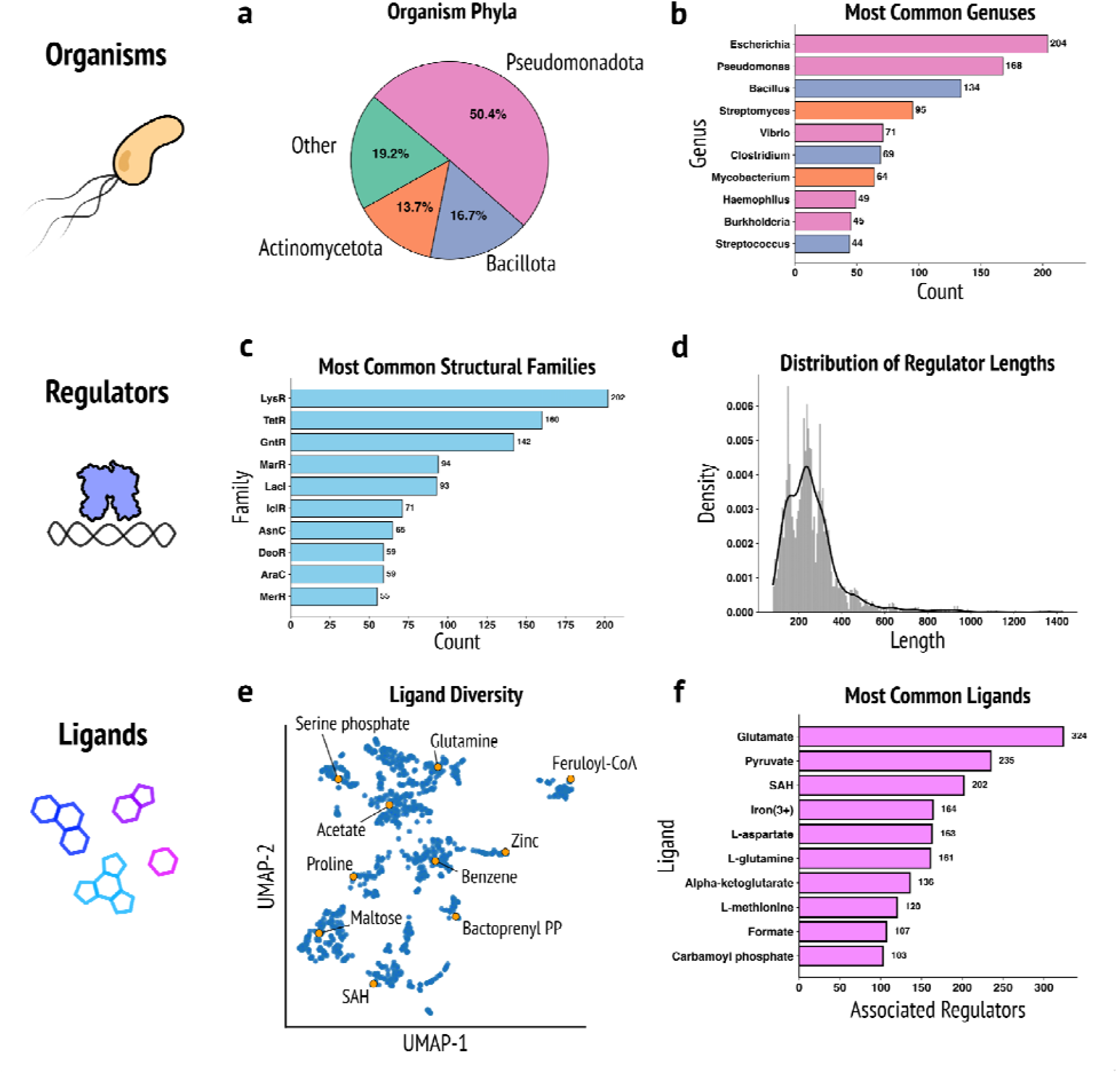
Organism, regulator, and ligand diversity of Ligify^DB^. (a) Distribution of organism phyla for regulators within the database. (b) The top 10 most common genuses. Bars are colored according to the organism’s phyla as shown in panel a. (c) The top 10 most common regulator structural families, indicated from regulator annotations. (d) Distribution of amino acid lengths among the 3,164 regulators in the database. (e) UMAP illustration of the structural diversity among the 1,362 ligands in the database. Highlighted chemicals are labeled and shown as orange dots. All other chemicals are shown as blue dots. (f) The top 10 most common ligands in the database.

### A web interface for biosensor search and visualization

The previous version of Ligify was built using Streamlit, which restricted data visualizations and accessibility. To provide an extensible interface where users can easily navigate and interact with Ligify^DB^ content, we created a dedicated web application using the React framework. Users can query the database via a search bar on the home page, which allows for chemical similarity search using a chemical SMILES input, or text search using a regulator’s RefSeq ID. Alternatively, users can browse the database from a data table on the “Browse” page, which allows for filtering entries based on sensor attributes, such as length or annotation, or filtering by full-text search.

The new React interface adds several new ways to visualize the database content (**Figure 3**). The 2D structures of all candidate ligands are rendered alongside an interactive AlphaFold model of the TF regulator protein. The protein sequence and predicted promoter sequence are displayed with length indicators. An interactive operon map allows users to study other proximally-encoded genes, which may provide additional insights into the true effector molecule recognized by the regulator. Metadata and literature for the associated enzyme is displayed, which is crucial for evaluating the quality of the prediction. A table containing the rank of the prediction, according to heuristics such as operon length and regulator-enzyme distance, is shown. Finally, a “plasmid builder” tool enables users to design a reporter circuit that uses the predicted regulator to control the expression of a fluorescent protein. The plasmid architecture is designed to be highly insulated and modular by default, incorporating strong terminators before and after each expression unit, and using modular bicistronic RBS elements to tunably control the expression of the regulator protein^23^. After selecting options for genetic elements, such as the fluorescent protein, the promoter and RBS strength driving the regulator, and the plasmid backbone, users can download an annotated genbank file of the designed plasmid.

**Figure 3:**
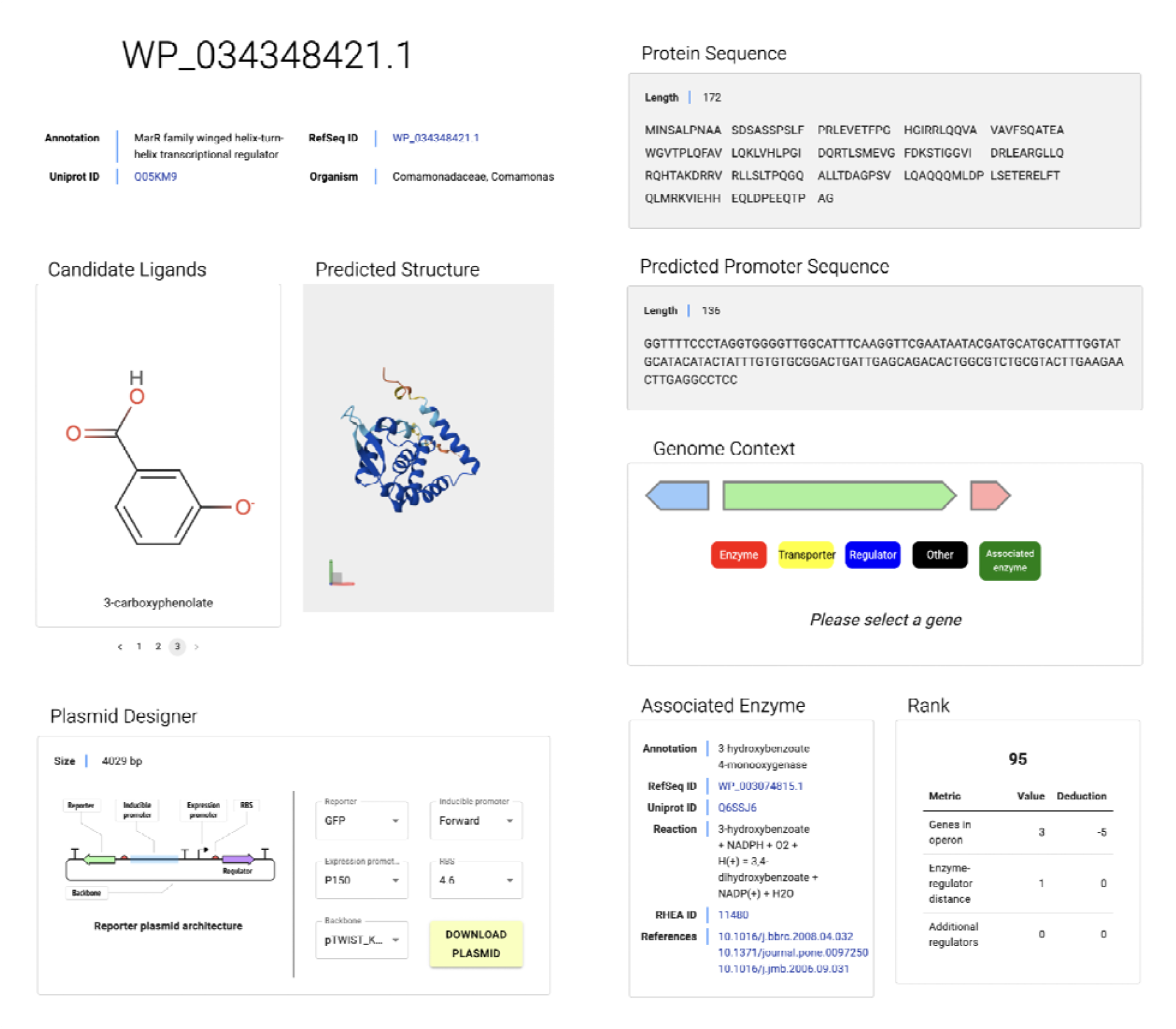
User interface of a representative Ligify^DB^ sensor page. Metadata and links to external databases are provided for each regulator, above their interactive AlphaFold structure and 2D structures of predicted candidate ligands. A plasmid designer tool enables users to create a reporter circuit using the regulator to control the expression of a fluorescent protein. The regulator sequence, predicted promoter sequence, interactive operon model, metadata on the associated enzyme, and calculated rank are also provided in the user interface.

A detailed documentation page provides context, statistics, a video tutorial, and guidance for users. The Ligify prediction model, workflow, and database creation pipeline are all described with illustrations, alongside a detailed comparison of Ligify^DB^ to the original version of Ligify. Descriptions of genetic elements in the “plasmid builder” tool are provided to help users understand which elements to tune and when. A contact form allows users to directly report any issues to the Ligify^DB^ administration team.

Finally, an option is available to download the entire 12 MB database, which should be useful for computational biologists who would benefit from programmatic access.

### Validation of biosensor predictions

We next sought to validate predictions made by Ligify 2.0. First, we used the groov^DB^ database, which contains experimentally validated genetic sensors, as a reference for benchmarking Ligify predictions^18^. Among the 216 sensors within groov^DB^ at the time of benchmarking, 69 were identified by Ligify while 147 were not (**Figure 4a**). This ∼32% predictive accuracy score is consistent with previous results, when Ligify was able to predict 31 of the 100 sensors within groov^DB^ nearly two years ago^14^.

**Figure 4:**
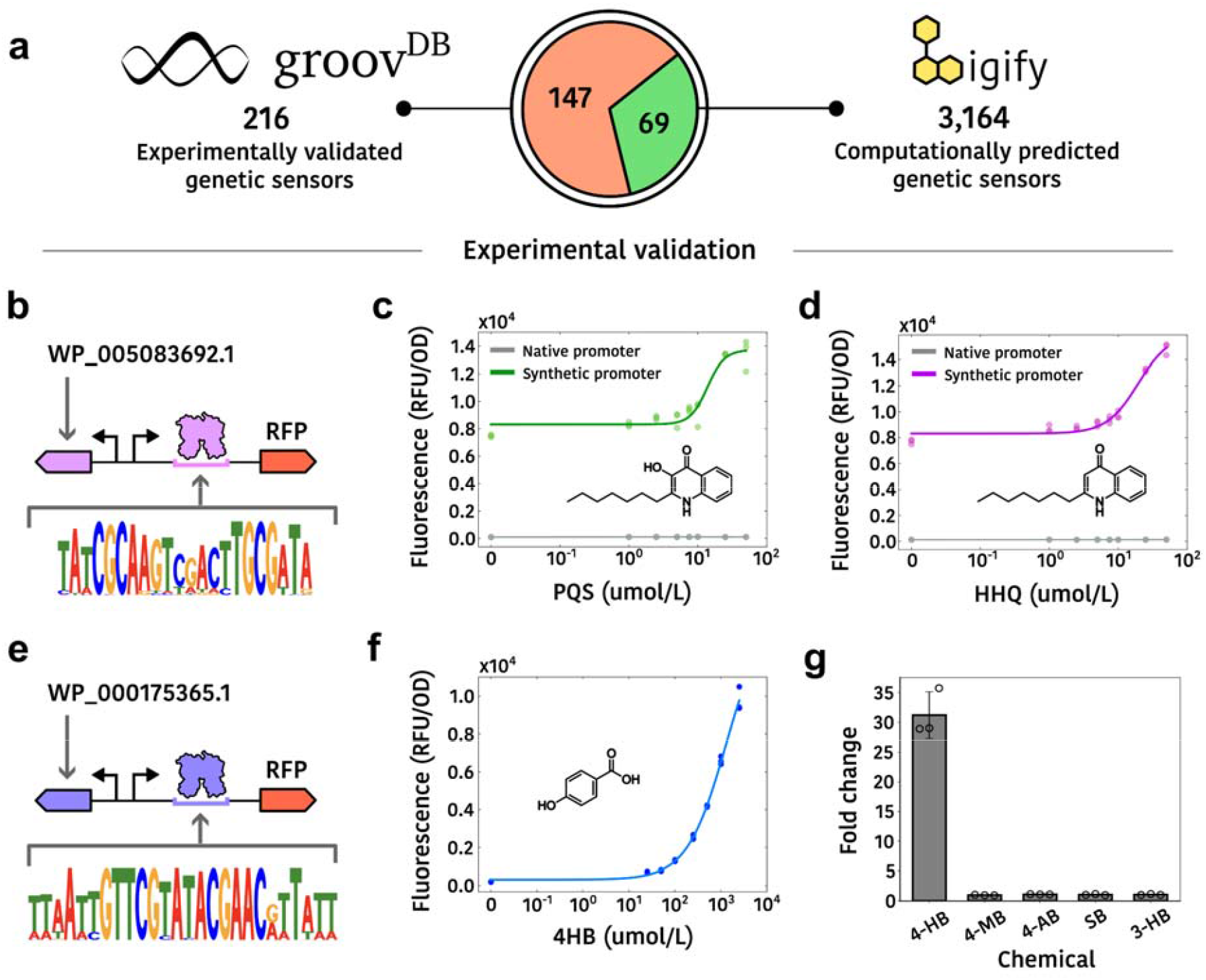
Validation of Ligify predictions. (a) Benchmarking Ligify predictions against experimentally validated sensors in groov^DB^. Among 216 sensors in groovDB, 69 were predicted by Ligify and 147 were not. (b-g) Experimental validation of sensors predicted by Ligify. (b) design of the PQS sensor’s genetic reporter circuit. The operator sequence was predicted by Snowprint. (c,d) Dose responses of the PQS sensor with PQS (2-heptyl-3-hydroxy-4-quinolone) and the PQS precursor HHQ (2-heptyl-4-quinolone). (e) design of the 4-hydroxybenzoate (4-HB) sensor’s genetic reporter circuit. (f) Dose response of the 4-HB sensor to 4-HB. (g) Response of the 4-HB sensor to 1 mM of 4-HB and various analogs. Error bars represent the standard deviation from the mean. All assays were performed in biological triplicate, with individual data points shown. Chemical names: 4-MB: 4-methylbenzoate, 4-AB: 4-aminobenzoate, SB: sodium benzoate, 3-HB: 3-hydroxybenzoate.

For further validation, we aimed to use Ligify to characterize transcription factors within pathogenic bacterial strains. Two regulators identified by Ligify were chosen for experimental testing, on the basis that they were from pathogens, they belonged to the easy-to-manipulate TetR family, their predicted ligands were commercially available, and they received a high confidence score, as determined by a set of previously described heuristics such as the number of genes within their operon. We first tested a sensor from *Mycobacterium abscessus* predicted to bind to the pseudomonas quinolone signal (PQS)^24^. Genetic circuits were constructed by expressing the sensor and separately expressing a red fluorescent protein, mScarlet-i3, using a synthetic promoter containing the PQS sensor’s operator sequence predicted by Snowprint^25^ (**Figure 4b**). This construct was able to respond about 2-fold to PQS and the precursor molecule HHQ (2-heptyl-4-quinolone) (**Figure 4c,d**). While suggested by a previous study, to our knowledge these results represent the first direct experimental evidence of a PQS-responsive TetR-family regulator in the pathogen *Mycobacterium abscessus*. To further validate Ligify predictions with another example, we built an analogous genetic circuit to measure the response of a regulator from the pathogen *Escherichia coli* O157:H7 predicted to respond to 4-hydroxybenzoate (4-HB) (**Figure 4e**). Consistent with *in vitro* characterizations performed by Roy & Ranjan, our genetic circuit produced a strong (up to 38-fold) and highly specific *in vivo* response to 4-HB^26^ (**Figure 4f,g**).

## Discussion

Here, we report a platform for predicted TF biosensors, comprising a database of 3,164 TFs associated with 1,362 ligands, and an accompanying interactive web interface for data navigation and visualization. Storing pre-computed prediction results in a static database delivers significant advantages over on-demand prediction tools, like TFBMiner and Ligify^13,14^. Static data is served much faster by requiring less computation and is easier to maintain by being less reliant on external resources, which would risk losing support over time. Access to all prediction results also enables an in-depth analysis of database content, as we provide here, and allows for downloading the entire database to facilitate incorporation into bioinformatic workflows.

The content in Ligify^DB^ was found to be relatively diverse, in terms of host phylogeny, TF structural families, and small molecule structures. While there is a bias towards model organisms, even the most represented organism (*E. coli*) comprised just 6% of the database. As expected, predicted ligands were largely composed of primary and secondary natural metabolites. Directed evolution approaches are likely necessary to yield TF biosensors responsive to synthetic molecules^27,28^. Our benchmarking results indicated that Ligify is able to faithfully predict true regulator-ligand interactions, and previous analyses suggest that predictive accuracy will increase as more data is deposited in the Rhea reaction database^14^. Finally, using Ligify, we were able to identify ligands for the previously characterized regulator HosA from *E. coli* O157:H7 and experimentally validate a PQS-responsive sensor from *Mycobacterium abscessus*.

We envision this resource as not just a static database, but a dynamic platform that will expand as enzyme reaction databases grow and new prediction methods are developed. For example, over 682 new chemicals have been added to the Rhea database since writing this manuscript. Other enzyme databases, such as KEGG and Brenda may also be used to increase the number of predicted TFs^29,30^. Beyond these expert curated databases, enzyme specificity prediction tools may be capable of expanding the scope of enzyme-ligand associations^31^. Separately, biophysical models may also prove useful for predicting TF-ligand interactions where the classic enzyme-association model fails. Machine learning models trained to predict TFs, such as DeepTFactor, can be used alongside AlphaFold to generate a large database of predicted TF structures^32^. Protein-ligand docking software can then be used to suggest possible ligands that fit the binding pocket of each predicted structure^33^. Finally, the resulting TF databases can be supplemented with complementary prediction tools, like Snowprint, to add information about hypothetical DNA binding sequences^34^. While only a tiny fraction of the several million sequenced TF biosensors have been characterized, combining biocuration with computational-guided experimental validation will likely accelerate progress towards mapping all prokaryotic TFs to the ligands they bind.

## Materials and Methods

### Database curation and analysis

All data processing was performed using Python v3.12.4. The following small molecules failed data fetching or were omitted due to poor likelihood of being true effector molecules (numbers indicate CHEBI IDs): water (15377), hydron (15378), dioxygen (15379), hydrogen peroxide (16240), nitrite (16301), carbon monoxide (17245), sulfite (17359), bicarbonate (17544), nitrate (17632), hydrosulfide (29919), ATP(4-) (30616), dTTP(4−) (37568), hydrogen phosphate (43474), Uridine triphosphase (46398), CoA (57287), FMNH2 (57618), UDP-α-D-glucuronate(3−) (58053). The following molecules were intentionally removed from the database due to their assumed role as cofactors rather than ligand effectors (numbers indicate occurrence): ATP (1102), Phosphate (837), ADP (836), Diphosphate (626), Ammonium (384), APAD (358), Carboxy-SAM (284), Coenzyme A (273), Oxygen (270), Acetyl CoA (209), 5-phosphoribosyl-PP (154), and D-glyceraldehyde 3-phosphate (130). All data is stored both in the public folder of the Ligify^DB^ source code, available on GitHub, as well as on Cloudflare R2, available for download.

### Building the Ligify^DB^ Web Application

The frontend web application was developed using React v18.3.1. Material UI (MUI) v7.2.0 was used as a user interface framework. Specific visualization components were built using a set of node libraries.

Chemical structures are displayed using smiles-drawer (v2.0.3), interactive protein structures are rendered using nightingale-structure (v5.6.1), and interactive operon diagrams are rendered using react-konva (18.2.10). Data is cached using Zustand (v5.0.8) to reduce subsequent page load times. Chemical similarity search is performed by converting the input SMILES code into a Morgan fingerprint using RDKit (radius: 2, size: 2048) and compared to fingerprints for all ligands within an AWS lambda function. The web application is deployed using AWS-Amplify.

### Benchmarking with groov^DB^

All sensors in the Ligify database were aligned to all sensors within the groov^DB^ database. Sensor entries from Ligify were considered matches with entries in groov^DB^ if their protein sequences were over 90% identical.

### Strains, plasmids, media, and chemicals

All biosensor circuits were characterized in *E. coli* DH10B (New England Biolabs). M9 media was used for routine cloning, fluorescence assays, unless specifically noted. M9 media were composed of the following filter-sterilized components: 33.9 g/L disodium phosphate, 15 g/L monopotassium phosphate, 2.5 g/L sodium chloride, 5 g/L ammonium chloride, 100 _μ_mol/L, CaCl2, 2 mmol/L MgSO4, 0.2% casamino acids, 0.4% glycerol, and 50 _μ_g/mL kanamycin. LB with 1.5% agar (BD) plates were used for routine cloning. The plasmids described in this work were constructed by Twist Biosciences. Ambeed was the vendor for the purchase of sodium 4-hydroxybenzoate (114-63-6) and PQS (108985-27-9).

### Fluorescence response assays

The following protocol was used to generate data shown in Figure 4c,d,f,g. Each biosensor plasmid was transformed into chemically competent *E. coli* cells. Three colonies were picked from each transformation and were grown overnight. The following day, 20 _μ_L of each culture was then used to inoculate separate wells in a 2 mL 96-deep-well plate (Corning, P-DW-20-C-S) sealed with an AeraSeal film (Excel Scientific) containing 900 _μ_L of M9 media. After 2 h of growth at 37 °C, cultures were induced with 100 _μ_L of M9 media containing either 10 _μ_L of the solvent (either water or DMSO) or 100 _μ_L of M9 media containing the target compound dissolved in the solvent (water or DMSO). Cultures were grown for an additional 4 h at 37 °C and 1000 r.p.m. and subsequently centrifuged (3500g, 4 °C, 10 min). The supernatant was removed, and cell pellets were resuspended in 1 mL PBS (137 mM NaCl, 2.7 mM KCl, 10 mM Na_2_HPO_4_, 1.8 mM KH_2_PO_4_, pH 7.4). One hundred microliters of the cell resuspension for each condition was transferred to a 96-well microtiter plate (Corning, 3904), from which the fluorescence (excitation, 569 nm; emission, 600 nm) and absorbance (600 nm) were measured using a plate reader (Biotek Neo2SM).

### Statistics and Reproducibility

All data in the text are displayed as mean ± standard deviation unless specifically indicated. All experimental assays were performed in biological triplicate, which represent three individual bacterial colonies picked from an agar plate. Bar graphs were all plotted in Python 3.10.6 using Matplotlib 3.8. Dose–response curves were plotted in Python 3.11.4 using Matplotlib 3.7.2.

## Data availability

All source code of Ligify^DB^ is open-sourced on GitHub under an MIT license and available here: https://github.com/groov-bio/ligify-ui/. The Ligify^DB^ website is publicly accessible here: https://ligify.groov.bio. This website is free and open to all users and there is no login requirement.

## Acknowledgements

We are grateful for helpful suggestions from Khalid K. Alam, Adam J. Meyer, and R. C. Baer.

## Author contributions

S.D.: Conceptualization, lead software development, and writing. N.Z. and P.T.: Experiments and data analysis. J.D.L. and J. J.: Assisted with software development, and helped revise the original draft. M. S. and P.A.S.: Supervision and writing. S.D.: Conceptualization, software contributions, data curation, and writing.

## Ethics declarations

S.D. has financial relationships with Retna Bio LLC. All other authors declare no competing interests.

